# *Ehrlichia* Notch signaling induction promotes XIAP stability and inhibits apoptosis

**DOI:** 10.1101/2023.01.06.523066

**Authors:** LaNisha L. Patterson, Caitlan D. Byerly, Regina Solomon, Nicholas Pittner, Duc Cuong Bui, Jignesh Patel, Jere W. McBride

## Abstract

*Ehrlichia chaffeensis* has evolved multiple strategies to evade innate defenses of the mononuclear phagocyte. Recently, we reported the *E. chaffeensis* TRP120 effector functions as a Notch ligand mimetic and a ubiquitin ligase that degrades the nuclear tumor suppressor, F-box and WD repeat domain-containing 7 (FBW7), a negative regulator of Notch. The Notch receptor intracellular domain (NICD) is known to inhibit apoptosis primarily by interacting with X-linked inhibitor of apoptosis protein (XIAP) to prevent degradation. In this study, we determined *E. chaffeensis* activation of Notch signaling increases XIAP levels, thereby inhibiting intrinsic apoptosis. Increased NICD and XIAP levels were detected during *E. chaffeensis* infection and after TRP120 Notch ligand mimetic peptide treatment. Conversely, XIAP levels were reduced in the presence of Notch inhibitor DAPT. Cytoplasmic colocalization of NICD and XIAP was observed during infection and a direct interaction was confirmed by co-immunoprecipitation. Procaspase levels increased temporally during infection, consistent with increased XIAP levels; however, knockdown of XIAP during infection significantly increased apoptosis and Caspase-3, −7 and −9 levels. Further, treatment with SM-164, a second mitochondrial activator of caspases (Smac/DIABLO) antagonist, resulted in decreased procaspase levels and increased caspase activation, induced apoptosis, and significantly decreased infection. In addition, iRNA knockdown of XIAP also decreased infection and significantly increased apoptosis. Moreover, ectopic expression of TRP120 HECT Ub ligase catalytically defective mutant in HeLa cells decreased NICD and XIAP levels and increased caspase activation compared to WT. This investigation reveals a mechanism whereby *E. chaffeensis* repurposes Notch signaling to stabilize XIAP and inhibit apoptosis.

**Author Summary:** *Ehrlichia chaffeensis* is a tick-borne, obligately intracellular bacterium that exhibits tropism for mononuclear phagocytes. *E. chaffeensis* survives by mobilizing various molecular strategies to promote cell survival, including modulation of apoptosis. This investigation reveals an *E. chaffeensis* initiated, Notch signaling regulated, antiapoptotic mechanism involving inhibitor of apoptosis proteins (IAPs). Herein, we demonstrate that *E. chaffeensis* induced Notch activation results in Notch intracellular domain stabilization of X-linked inhibitor of apoptosis protein (XIAP) to inhibit intrinsic apoptosis. This study highlights a novel mechanistic strategy whereby intracellular pathogens repurpose evolutionarily conserved eukaryotic signaling pathways to engage an antiapoptotic program for intracellular survival.

## Introduction

*Ehrlichia chaffeensis* is an obligately intracellular Gram-negative bacterium, and the etiologic agent of human monocytotropic ehrlichiosis (HME), a life-threatening emerging tick-borne zoonosis (1). *E. chaffeensis* preferentially infects mononuclear phagocytes and has evolved sophisticated molecular-based strategies to evade host defense mechanisms for survival (2–18). *E. chaffeensis* immune evasion strategies are mediated, in part, by tandem repeat proteins (TRPs). TRPs are type 1 secretion system (T1SS) effectors that also elicit strong host antibody responses during infection (19–22). Notably, TRP120 decorates the surface of dense-cored ehrlichiae and has multiple moonlighting functions including as a transcription factor, HECT E3 ubiquitin ligase, and cellular signaling ligand mimetic to repurpose host cell signaling (2, 9, 13–16, 18, 23). These TRP120 functions directly impact host gene expression and chromatin epigenetics, pathogen-host interactions, and cellular signaling (2, 4–7, 9, 12–14, 18, 24, 25).

Two apoptosis pathways, extrinsic and intrinsic, have been defined and are well characterized. The extrinsic pathway is activated through a death ligand receptor resulting in the activation of Caspase-8 and induction of the execution pathway leading to apoptosis (26–28). By comparison, the intrinsic pathway is initiated by various non-receptor mediated stimuli that result in mitochondrial changes, specifically mitochondrial permeability transition (MPT). MPT results in cytochrome c release, triggering formation of a complex known as an apoptosome and subsequent Caspase-9 activation resulting in apoptosis (29, 30). Execution of apoptosis occurs when Caspase-8 and/or −9 cleave inactivated executioner Caspase-3/7 into their active forms, leading to the cleavage of various downstream targets important for cell survival (29, 31, 32). Modulation of several proteins that control and regulate apoptotic mitochondrial events (intrinsic apoptosis) occur during *E. chaffeensis* infection including Bcl-2, BirC3, and others have demonstrated downregulation of apoptotic inducers, such as Bik, BNIP3L, and hematopoietic cell kinase (HCK) (33). The *E. chaffeensis* effector, ECH0825, is also known to suppress apoptosis by inhibiting Bax-induced apoptosis by increasing mitochondrial manganese superoxide dismutase (MnSOD) to reduce reactive oxygen species-mediated damage (25). Although the manipulation of intrinsic apoptosis as a survival mechanism for *E. chaffeensis* has been previously reported, there remain significant unanswered questions about the mechanisms involved.

Our laboratory has recently reported that *E. chaffeensis* evasion of macrophage host defenses involves activation of conserved host signaling pathways, including Wnt, Notch and Hedgehog (2, 13, 14). Notably, TRP120 activates the evolutionarily conserved Notch signaling pathway using a novel molecularly defined pathogen-encoded Notch SLiM ligand mimetic found within the tandem repeat (TR) domain (13). Notch signaling plays significant roles in cellular homeostasis, MHC Class II expansion, B- and T-cell development, and modulation of innate immune mechanisms such as autophagy and apoptosis (34–39). Recently, we have reported *E. chaffeensis* TRP120-induced Notch signaling results in downregulation of toll-like receptor (TLR) 2/4 expression (5). Moreover, we determined that TRP120 degrades the Notch negative regulator, FBW7, resulting in increased levels of several oncoproteins, including the Notch intracellular domain (NICD), which regulates cell survival and apoptosis (15). Therefore, *E. chaffeensis* induced Notch signaling and increased levels of NICD during *E. chaffeensis* infection may play an important role in inhibiting apoptosis.

Caspases are the enzymes primarily responsible for mediating apoptosis (26, 29, 40), and apoptosis can be blocked by inhibiting caspase activity. The X-linked inhibitor of apoptosis protein (XIAP) is the most potent inhibitor of apoptosis (IAP) (41, 42). XIAP directly binds and inhibits initiator and executioner caspases, including Caspases-9 and −3, respectively (42–46). Interestingly, Lui, et al demonstrated that NICD suppresses host cell apoptosis by increasing XIAP stability (47). NICD and XIAP interaction prevents ubiquitination and degradation of XIAP, thereby inhibiting apoptosis. Moreover, Caspase-3 and −8 are known to cleave XIAP into two fragments (BIR1-2 and BIR3-RING), leading to differential inhibition of extrinsic and intrinsic apoptotic pathways (48). BIR3-RING fragments are potent inhibitors of Caspase-9, resulting in inhibition of Bax-mediated (intrinsic) apoptosis. Therefore, Notch activation and FBW7 degradation during *E. chaffeensis* infection may stabilize XIAP as a mechanism to inhibit intrinsic, caspase-dependent apoptosis.

In this study, we reveal a novel mechanism whereby *E. chaffeensis* inhibits intrinsic apoptosis through Notch activation resulting in NICD stabilization of XIAP. Inhibition of apoptosis through modulation of Notch signaling provides further evidence that *E. chaffeensis* hijacks evolutionarily conserved signaling pathways primarily to evade innate host defense mechanisms.

## Results

### *E. chaffeensis* infection and TRP120 increases XIAP levels

We recently demonstrated that NICD levels temporally increase during *E. chaffeensis* infection (15). Increased levels of NICD were associated with TRP120 ubiquitination and degradation of a Notch negative regulator, FBW7. NICD is known to directly bind the XIAP BIR-RING domain prevent XIAP autoubiquitination and degradation (47), thereby inhibiting apoptosis. To investigate potential XIAP upregulation during infection, THP-1 cells were incubated with *E. chaffeensis* (MOI 50), and XIAP protein and transcription levels were analyzed by immunoblot and qPCR. Levels of XIAP were unchanged in uninfected THP-1 cells (Fig. 1A). In comparison, increases in both NICD and XIAP protein levels were demonstrated over the course of infection, with significant increases detected at 24, 48 and 72 hpi (Fig. 1A). Interestingly, additional cleaved fragment was observed at 48 and 72 hpi and identified as the BIR3-RING domain of XIAP (Fig. 1A). When cleaved, BIR3-RING also acts as a potent inhibitor of the intrinsic apoptotic pathway by binding the Caspase-9 monomer preventing its cleavage and heterodimerization. Moreover, transcriptional levels of XIAP were also shown to be significantly upregulated in a temporal manner (Fig. 1B). XIAP expression was determined in THP-1 cells treated with recombinant TRP120 tandem repeat domain (rTRP120-TR), or the recently described TRP120 Notch ligand memetic SLiM (TRP120-TR-P6) peptide. A significant temporal increase in XIAP levels was detected in cells treated with rTRP120-TR (Fig. 1C) or TRP120-TR-P6 peptide (Fig. 1D), demonstrating *E. chaffeensis* TRP120 promotes increased XIAP levels.

**Fig. 1.**
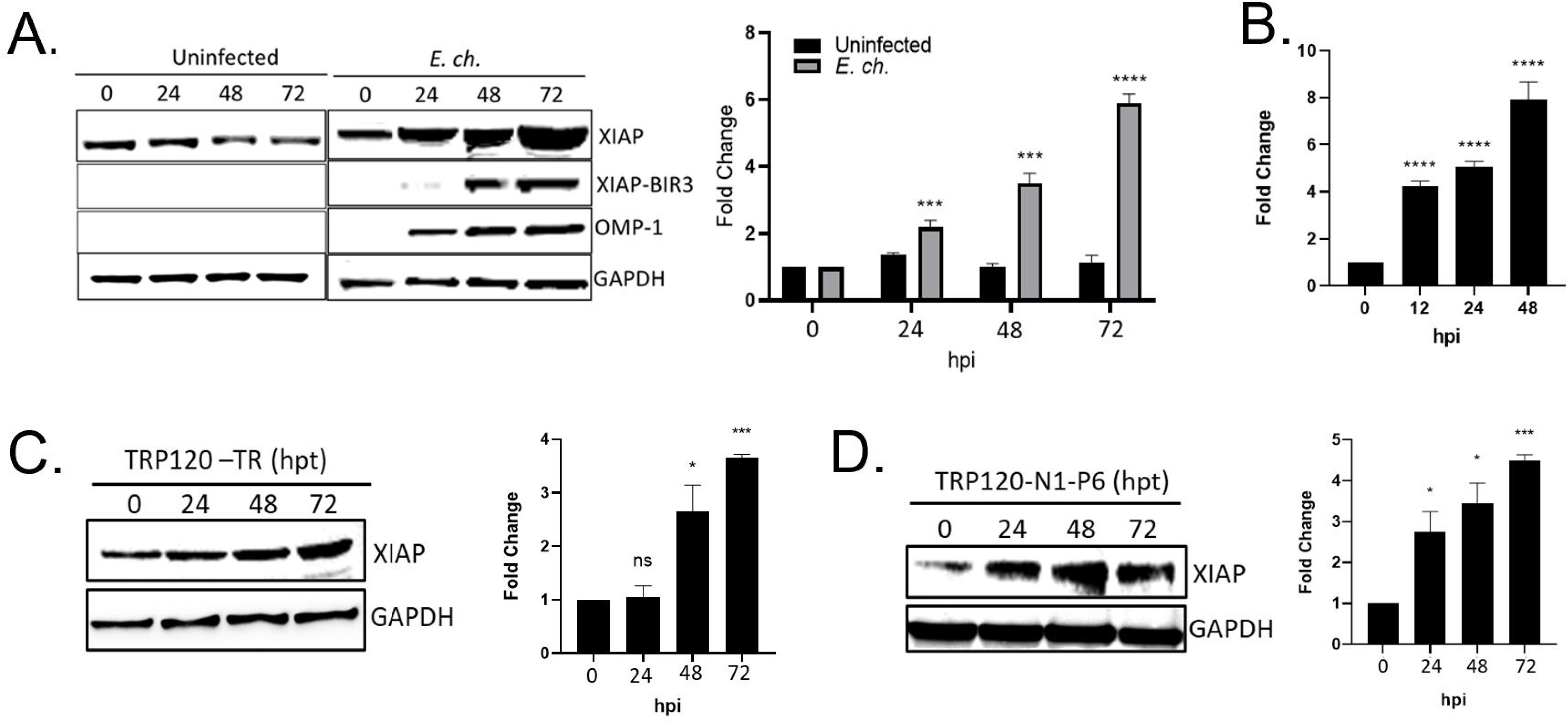
XIAP levels increase during *E. chaffeensis* infection. (A) Immunoblot of XIAP, cleaved XIAP-Bir3 and OMP-1 expression in uninfected and *E. chaffeensis-infected* THP-1 cells at 0-72 hpi (MOI 50). The major outer membrane protein-1 (OMP-1) is used to confirm infection. GAPDH was utilized as a loading control. Fold differences in XIAP levels in *E. chaffeensis-infected* and uninfected THP-1 cells. (B) Changes in XIAP transcript levels in *E. chaffeensis-infected* and uninfected THP-1 cells (0 hpi) as determined by RT–qPCR analysis. (C) Immunoblot and quantification of XIAP in TRP120-TR-treated THP-1 cells. (D) Immunoblot and quantification of XIAP in TRP120-N1-P6 memetic peptide-treated THP-1 cells. Bar graphs represent mean ± SD. ****, P < 0.0001. Experiments were performed in triplicate (n=3) and representative images are shown.

### NICD interacts with XIAP during infection

To determine if NICD was directly binding XIAP, confocal microscopy was performed to visualize XIAP/NICD colocalization. Interestingly, strong NICD and XIAP colocalization according to Mander’s coefficient (MC) was observed in *E. chaffeensis-infected* cells at 24 (MC = 0.9), 48 (MC = 0.9) and 72 (MC = 0.8) hpi (Fig. 2A). XIAP levels were also temporally increased at 24, 48 and 72 hpi based on mean fluorescence intensity (Fig. 2B). By immunoblot, significant temporal increases in XIAP and NICD were detected (Fig. 2C). Co-immunoprecipitation of XIAP and NICD from uninfected and *E. chaffeensis-infected* lysates (24 hpi) demonstrated direct interaction and increased XIAP and NICD levels (Fig. 2D).

**Fig. 2.**
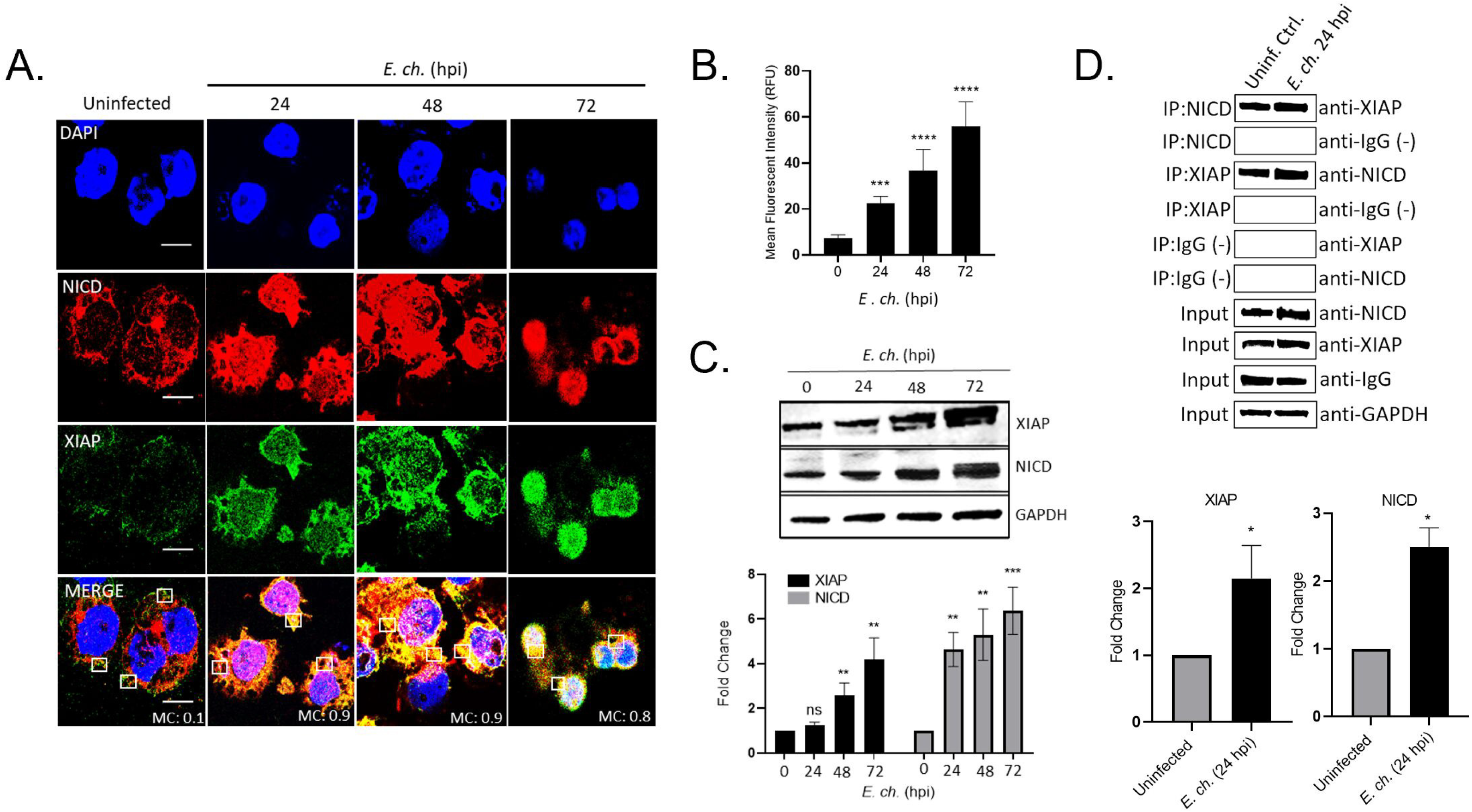
NICD interaction with XIAP during *E. chaffeensis* infection. (A) Uninfected or *E. chaffeensis-infected* THP-1 cells at 24, 48 and 72 hpi (MOI 50) probed for endogenous NICD (red) or endogenous XIAP (green) demonstrate colocalization by immunofluorescent confocal microscopy (scale bar = 10 μm). Colocalization was quantitated by Mander’s coefficient (0 no colocalization; +1 strong colocalization). Mean fluorescence intensity of the XIAP protein expression in 20 THP-1 cells. Mean pixel values were obtained using the ImageJ Measure Analysis tool. Background intensity was determined and subtracted from the fluorescence intensity value of the cells. The mean value from each group was an average of 20 cells. (C) Immunoblot and fold differences of XIAP or NICD in *E. chaffeensis*-infected cells at 0-72 hpi (MOI 50). GAPDH was utilized as a loading control. (D) Co-IP and reverse Co-IP demonstrate the direct interaction between XIAP and NICD at 24 hpi compared to the IgG negative control. Western blot analysis was normalized to GAPDH expression. Quantification of NICD or XIAP levels from one representative CO-IP experiment are shown. Experiments were performed in triplicate (n=3) and representative images are shown.

### Notch activation and XIAP stabilization by NICD

To confirm that the increase in XIAP levels was a direct result of Notch activation, THP-1 cells were pre-treated with DAPT, a Notch γ-secretase inhibitor. Cells pre-treated with DAPT inhibitor and infected with *E. chaffeensis* (MOI 50) exhibited decreased XIAP levels 24 and 48 hpi (Fig. 3A). Similar results were also shown with SAHM1 treatment, which prevents assembly of the active Notch transcriptional complex (Fig. S1). To further determine the direct relationship between NICD and XIAP levels during *E. chaffeensis* infection, uninfected and *E. chaffeensis-infected* THP-1 cells were treated at 1 hpi with DAPT and SM-164, a second mitochondrial activator of caspases (Smac/DIABLO) mimetic compound that antagonizes inhibitor of apoptosis proteins (IAPs) that promotes activation of caspases and apoptosis (45). Cell death was induced by TNF-α, followed by subsequent infection with *E. chaffeensis* (MOI 50). H&E staining of THP-1 cells treated with DAPT/SM-164 did not contain morulae, displayed significant cell death, and had significantly decreased ehrlichial load compared to uninfected, *E. chaffeensis-infected,* DMSO alone, or with DAPT or SM-164 alone treated cells (Figs. 3B and C).

**Fig. 3.**
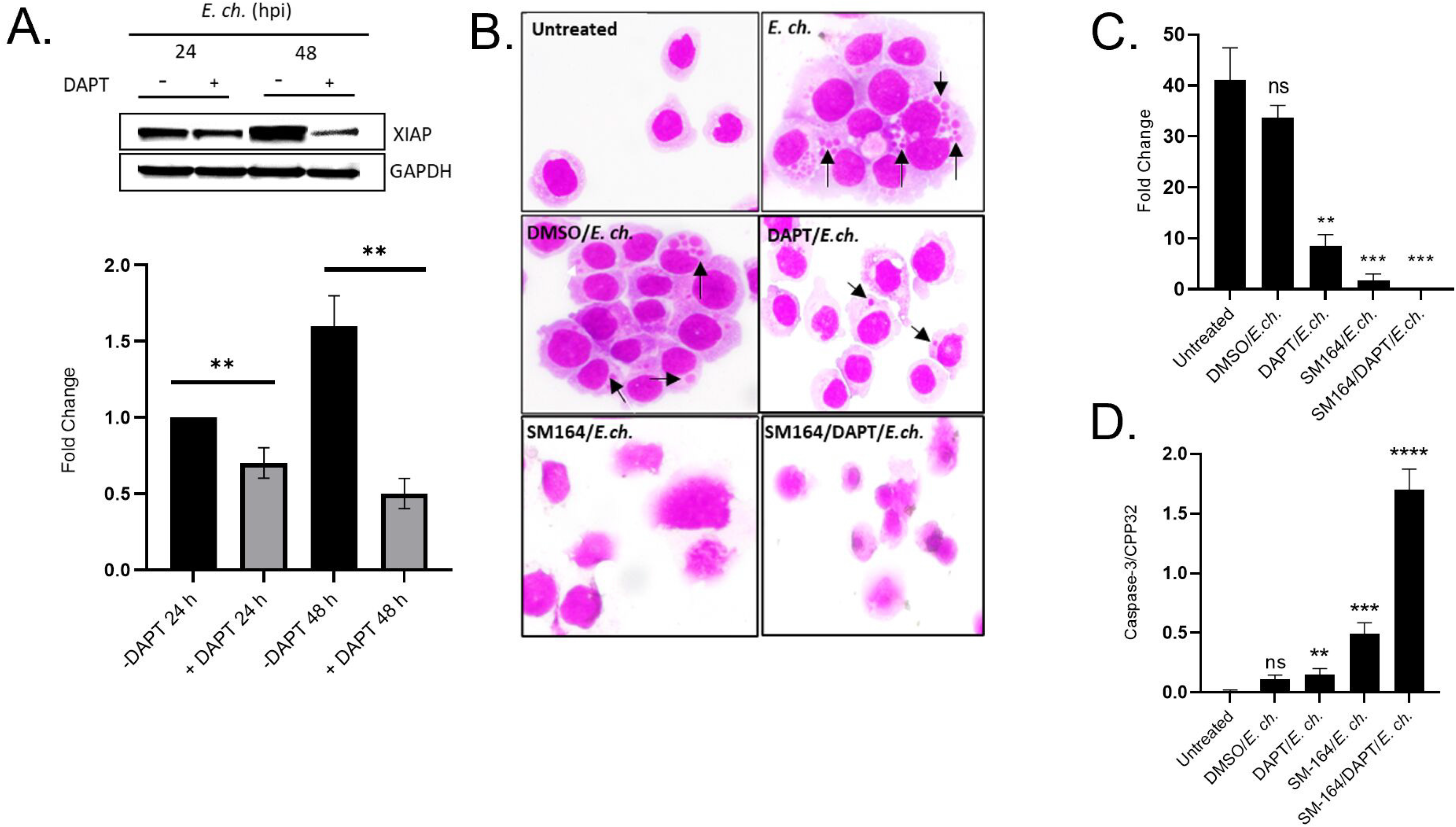
Notch activation stabilizes XIAP during *E. chaffeensis* infection. (A) Immunoblot and fold differences of XIAP expression in *E. chaffeensis-infected* THP-1 cells with or without DAPT pre-treatment at 24 or 48 hpi. (B) H&E stained uninfected or *E.chaffeensis-infected* THP-1 cells untreated or pre-treated with DMSO or SM164 (12 h) alone or in combination with DAPT (1 h). Cell death was stimulated by TNF-α prior to the addition of *E. chaffeensis* (MOI 50). Black arrows identify *E. chaffeensis* morulae. (C) Fold-change difference in *dsb* transcript levels in cells depicited in Fig. 3B. (D) Quantification of Caspase-3/CPP32 activity determined at an absorbance of 405 nm. Bar graphs represent mean ± SD. ****, P < 0.0001; ns = not significant. Experiments were performed in triplicate (n=3) and representative images are shown.

To further confirm Notch signaling promotes cell survival and ehrlichial infection, Caspase-3/CPP32 activity was determined by measuring the absorbance of DEVD-pNA, a Caspase-3 substrate (49). An approximately 2-fold increase in Caspase-3/CPP32 levels were detected in THP-1 cells treated with both DAPT and SM-164 compared to uninfected, DMSO treated, or *E. chaffeensis* infected treated with either DAPT or SM-164 (Fig 3D). Significant upregulation of Caspase-3/CPP32 activity was also demonstrated in the SM-164 treated compared to untreated cells; however, DAPT/SM-164 combination treatment demonstrated significantly higher Caspase-3/CPP32 levels (Fig 3D). Collectively, these data suggest that XIAP is stabilized by NICD during *E. chaffeensis* infection.

### NICD stabilization of XIAP results in antiapoptotic activity and increased infection

The effect of XIAP on *E. chaffeensis* infection was examined using siRNA knockdown (KD) of XIAP in THP-1 cells. siRNA KD of XIAP resulted in a 77% KD efficiency (Fig. 4A). *E. chaffeensis* infection was significantly decreased 24 hpi in XIAP-KD cells compared to scrambled control siRNA-treated cells (Figs. 4B and C). To determine if the decrease in ehrlichial load was caused by the induction of apoptosis due to XIAP destabilization, cell viability was determined by flow cytometry with the Muse Count & Viability Kit. XIAP-KD cells exhibited a viability of ~8% compared to 81% in scr-KD cells (Fig. 4D). Furthermore, XIAP-KD cells displayed morphological changes associated with apoptosis including shrinkage of the cell, fragmentation into membrane-bound apoptotic bodies and nuclear fragmentation (Fig. 4D).

**Fig. 4.**
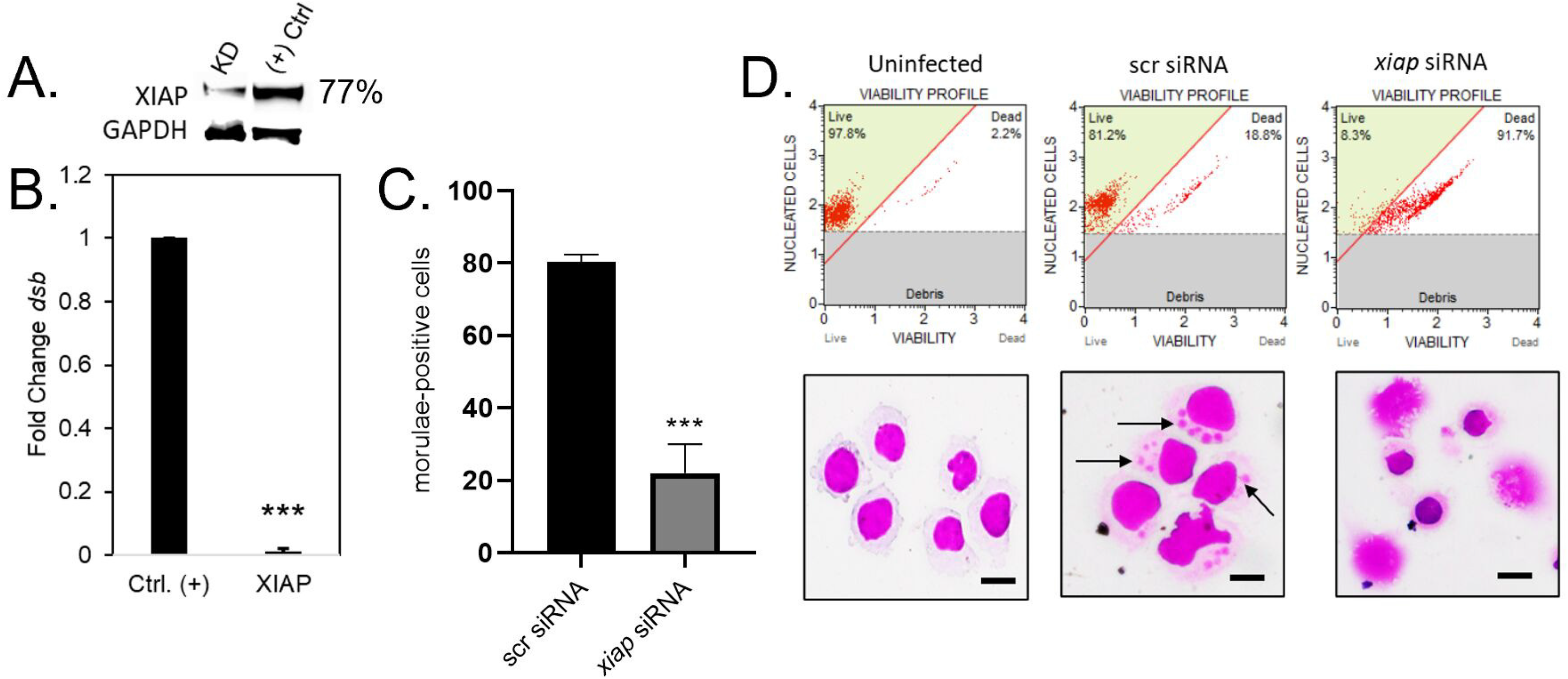
XIAP enhances cell viability to promote *E. chaffeensis* infection. (A) Immunoblot depicting knockdown efficiency of *XIAP* in siRNA knockdown cells compared to positive control from *E. chaffeensis* THP-1 cells harvested at 24 hpi. Percent knockdown of XIAP relative to positive control is shown to the right of the immunoblot. GAPDH is utilized as a loading control. (B) Small interfering RNA-transfected (siRNA) THP-1 cells were infected with *E. chaffeensis* (MOI 100, 24 hpt). Scrambled siRNA (scrRNA) was transfected as an infected positive control. *E. chaffeensis* infection was quantified as fold change at 24 hpi and was determined by qPCR amplification of the *dsb* gene. Knockdowns were performed with at least three biological and technical replicates for *t*-test analysis. (C) Quantification of morulae positive cells in scr or *XIAP* siRNA-treated THP-1 cells by H&E staining. (D) Representative image of H&E stained uninfected or *E. chaffeensis-* infected scr or *XIAP* siRNA-treated cells (arrows identify *E. chaffeensis* morulae). Quantitative analysis of cell viability by the Muse^®^ Count & Viability Kit for uninfected or *E. chaffeensis-infected* scr or *XIAP* siRNA-treated cells is shown. Bar graphs represent mean ± SD. ****, P < 0.001. Experiments were performed in triplicate (n=3) and representative images are shown.

### XIAP stabilizes pro-caspase levels to inhibit apoptosis

Smac/DIABLO is a cytosolic antagonist of IAPs (50). To determine the significance of IAPs during *E. chaffeensis* infection, THP-1 cells were treated with SM-164, a Smac/DIABLO mimetic compound. Cell death was induced by TNF-α, followed by subsequent infection with *E. chaffeensis.* Treatment with SM-164 resulted in a significant reduction in ehrlichial infection as determined by confocal microscopy and qPCR of the *dsb* gene (Figs. 5A and B). Importantly, cells treated with SM-164/TNF-α had 10-20% viability and exhibited morphological changes of apoptosis including membrane blebbing, nuclear fragmentation and cell shrinkage (Figs. 5A and C, S2A). In comparison, untreated or DMSO-treated cells had cell viabilities ranging from 82-92% and 63-70%, respectively (Figs. 5C, S2A). Additionally, there were unremarkable morphological changes associated with untreated or DMSO-treated cells (Fig. 5A). These data further suggests that NICD stabilization of XIAP results in antiapoptotic activity during *E. chaffeensis* infection. In addition, an increase in early and late apoptotic cells were shown in SM-164/TNF-α-treated cells compared to untreated, DMSO or either SM-164 or DMSO/TNF-α treated cells. Importantly, pre-incubation of THP-1 cells with Caspase-9 inhibitor, Z-LEHD-FMK TFA, prior to SM-164/TNF-α treatment reversed apoptotic effects demonstrated with SM-164/TNF-α (Figs. 5A-C, S2A-B). Collectively, these data demonstrate increases in XIAP levels inhibit intrinsic apoptosis during *E. chaffeensis* infection.

**Fig. 5.**
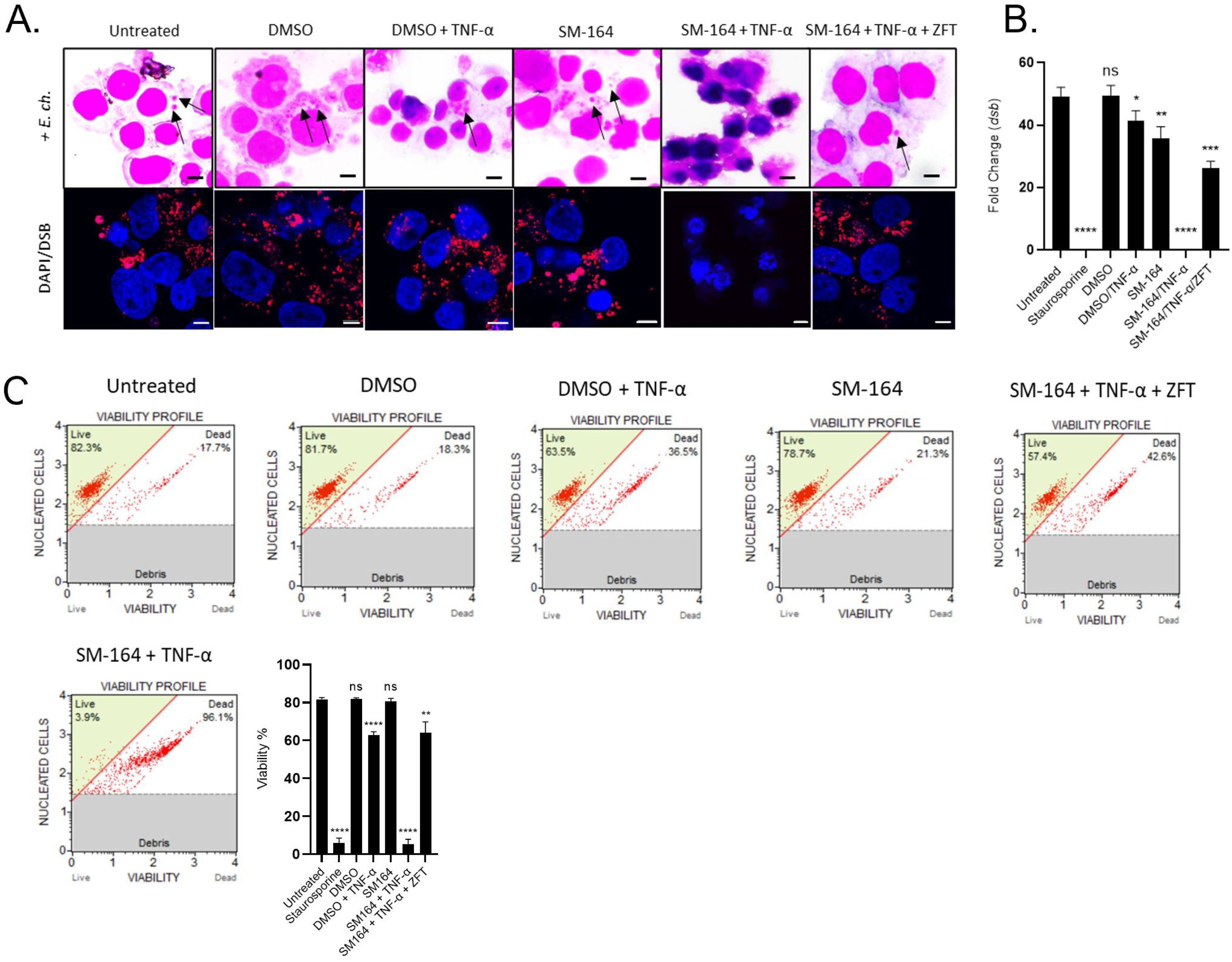
XIAP stabilizes pro-caspase levels to inhibit apoptosis. *E. chaffeensis-infected* THP-1 cells (MOI 50) were pre-treated with DMSO or SM-164 alone (100 nM, 12 h) or SM-164 (100 nM, 12 h) in combination with Caspase-9 inhibitor, Z-LEHD-FMK TFA (20 μM, 2 h). Cell death was stimulated with TNF-α (100 ng/ml) or staurosporine (100 ng/ml, positive apoptosis control). (A) The presence of *E. chaffeensis* determined by H&E staining (black arrows) and immunofluorescent confocal microscopy. Infected THP-1 cells were probed with anti-DSB and tetramethylrhodamine isothiocyanate (TRITC) conjugate (red) to confirm the presence of morulae by immunofluorescent confocal microscopy. Nuclei were stained with 4’,6’-diamidino-2-phenylindole DAPI (blue). Apoptotic cells were identified by visualization of nuclear morphology by DAPI (bar = 10μm). (B) Fold change difference of *dsb* transcript levels in the indicated cell samples. (C) Cell viability was determined using the Muse^®^ Count & Viability Kit and quantification of cell viability percentage in the previously mentioned treatment groups is shown. Bar graphs represent mean ± SD. ****, P < 0.0001; ns = not significant. Experiments were performed in triplicate (n=3) and representative images are shown.

### Pro-Caspase levels during *E. chaffeensis* infection

Studies have demonstrated that XIAP differentially inhibits caspases through its baculovirus IAP repeat (BIR) domains (48). The BIR2 domain directly binds apoptotic executioner Caspase-3 and −7 using a two-site interaction mechanism for inhibition of apoptosis (51). XIAP also sequesters Caspase-9 in a monomeric state using the BIR3 domain, preventing the catalytic activity of Caspase-9 (45). Therefore, XIAP is directly associated with inhibition of downstream caspases. During *E. chaffeensis* infection, temporal levels of Caspase-3, −7 and −9 increased during infection (Fig. 6A). Temporal increases in Caspase-3, −7 and −9 gene transcription was also detected during *E. chaffeensis* infection (Fig. S3).

**Fig. 6.**
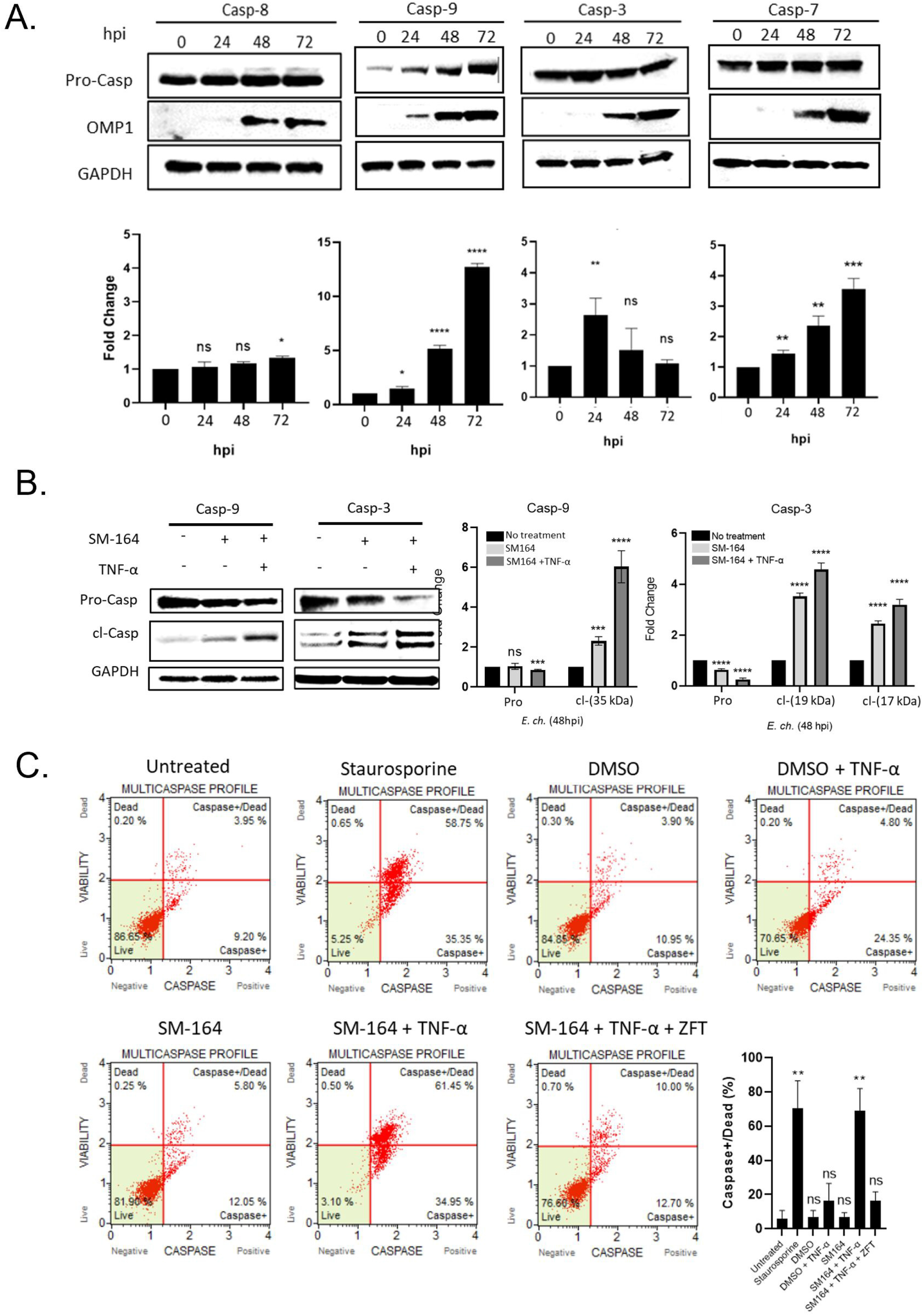
XIAP stabilizes pro-caspase levels to inhibit apoptosis during *E. chaffeensis* infection. (A) Immunoblots and fold change in pro-Casp-8, −9, −3 and −7 in *E. chaffeensis*-infected cells at 0, 24, 48 and 72 hpi. Outer membrane protein 1 (OMP-1) is a major immunodominant protein used to confirm infection. GAPDH is utilized as a loading control. (B) Immunoblot and quantification of the fold change of *E. chaffeensis*-infected cells untreated or pre-treated with SM-164 alone or in combination with TNF-α. Cells were probed for total and cleaved Caspase-3 or 9 and fold change differences are shown. (C) *E. chaffeensis*-infected THP-1 cells were pre-treated with DMSO or SM-164 alone (100nM, 12h) or SM-164 (100nM, 12h) in combination with Caspase-9 inhibitor, Z-LEHD-FMK TFA (20μM, 2 h). Cell death was stimulated with TNF-α (100 ng/ml) or staurosporine (100 ng/ml, positive apoptosis control) in the indicated samples. Percentages of live, caspase+, caspase+ and dead, total caspase+, and dead cells determined using the Muse^®^ MultiCaspase Kit. Quantification of Caspase+/dead cells (%) is shown. Bar graphs represent mean ± SD. **, P < 0.01; ns = not significant. Experiments were performed in triplicate (n=3) and representative images are shown.

To demonstrate that XIAP was directly associated with downstream caspase inhibition, THP-1 cells were treated with SM-164 and TNF-α, infected with *E. chaffeensis* infection (MOI 50) and Caspase-9 and −3 levels were determined. The SM-164/TNF-α treated cells had significantly lower levels of pro-Caspase-9 and −3, and significantly increased levels of cleaved (active) Caspase-9 and −3 (Fig. 6B). To demonstrate that XIAP was directly associated with downstream caspase inhibition, THP-1 cells were treated with SM-164 in combination with TNF-α, infected with *E. chaffeensis* infection (MOI 50) and multi-caspase levels were determined by flow cytometry. A significant increase in the percentage of caspase+/dead cells with SM-164/TNF-α treatment was detected (Fig. 6C). These data demonstrate increased XIAP levels inhibit caspase activation and apoptosis during *E. chaffeensis* infection.

### FBW7 ubiquitination and proteasomal degradation stabilizes XIAP

We have recently demonstrated FBW7 degradation during *E. chaffeensis* infection results in increased levels of FBW7 regulated oncoproteins including NICD, phosphorylated c-Jun, MCL-1 and cMYC (15). Data demonstrated that TRP120 ubiquitination of FBW7 results in FBW7 degradation, enhancing infection. To determine the effect of FBW7 and NICD on XIAP and Caspase-3 and −9 activation, HeLa cells were transfected with catalytically-inactive TRP120 (TRP120-C520S) and treated with TRP120-N1-P6 Notch memetic peptide for 2 h to activate Notch signaling followed by induction of cell death by TNF-α. HeLa cells ectopically expressing TRP120-C520S displayed increased cleaved Caspases-3 and −9 compared to uninfected or TRP120-WT transfected cells (Fig. 7). Collectively, these data demonstrate that TRP120 FBW7 degradation stabilizes NICD for subsequent increased XIAP expression and inhibition of caspase activation during *E. chaffeensis* infection.

**Fig. 7.**
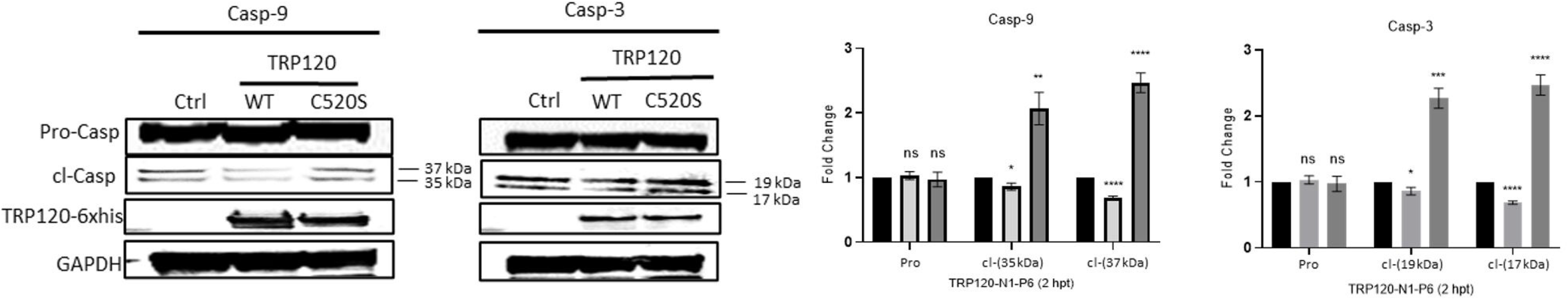
FBW7 degradation during *E.ch.* Infection enhances XIAP stability and caspase activation. HeLa cells were transfected with empty vector, TRP120-WT or catalytic-inactive TRP120 (TRP120-C520S) for 24 h, stimulated with TRP120-N1-P6 memetic peptide to activate Notch signaling (2 h) and cell death was induced by TNF-α (100 ng/ml, 48 h). (A) Immunoblot of Pro- and cleaved Caspase-3 or −9 in vector, TRP120-WT or TRP120-C520S-treated cells. Fold change in Pro- and cleaved Caspase-3 or −9 levels in TRP120-WT or TRP120-C520S-treated cells compared with empty vector are shown. Bar graphs represent mean ± SD. ****, P < 0.0001. Experiments were performed in triplicate (n=3) and representative images are shown.

## Discussion

Inhibition of host cell apoptosis is an important survival strategy utilized by *E. chaffeensis* (33, 52, 53). Previous studies have demonstrated the T4SS effector protein, ECH0825, inhibits host cell apoptosis in human monocytes (54). ECH0825 localizes to mitochondria and inhibits Bax-induced apoptosis by increasing mitochondrial manganese superoxide dismutase (MnSOD) and reducing reactive oxygen species-mediated damage (54). Further, upregulation of apoptotic inhibitor genes during *E. chaffeensis* infection, including BCL-2 and BIRC3, and downregulation of apoptotic inducers, such as BIK, BNIP3L, and hematopoietic cell kinase have also been reported (33). However, there is little information related to *E. chaffeensis* modulation of caspase activation. In this study, we have identified a mechanism by which *E. chaffeensis* TRP120 effector activates the Notch signaling pathway to inhibit caspase-dependent apoptosis through NICD and XIAP interaction.

We recently identified a TRP120 Notch SLiM ligand memetic motif responsible for Notch activation during *E. chaffeensis* infection (13). Additionally, we demonstrated that *E. chaffeensis* and rTRP120 activate Notch signaling to downregulate TLR 2/4 expression for intracellular survival (5). TRP120 Notch activation was recently confirmed to occur through a TRP120 tandem repeat (TRP120-TR) Notch SLiM ligand memetic that directly binds to the Notch receptor in a region containing the known ligand binding region (13). Although a Notch SLiM memetic has been recently identified, its role in *E. chaffeensis* survival has not been fully elucidated. In this study, we investigated the functional implications of TRP120 Notch SLiM memetic activation during *E. chaffeensis* infection and defined a novel antiapoptotic mechanism involving inhibitor of apoptosis (IAP) proteins potently inhibiting the catalytic activity of caspases through regulation of Notch signaling in the illustration abstract (Fig. 8).

**Fig. 8.**
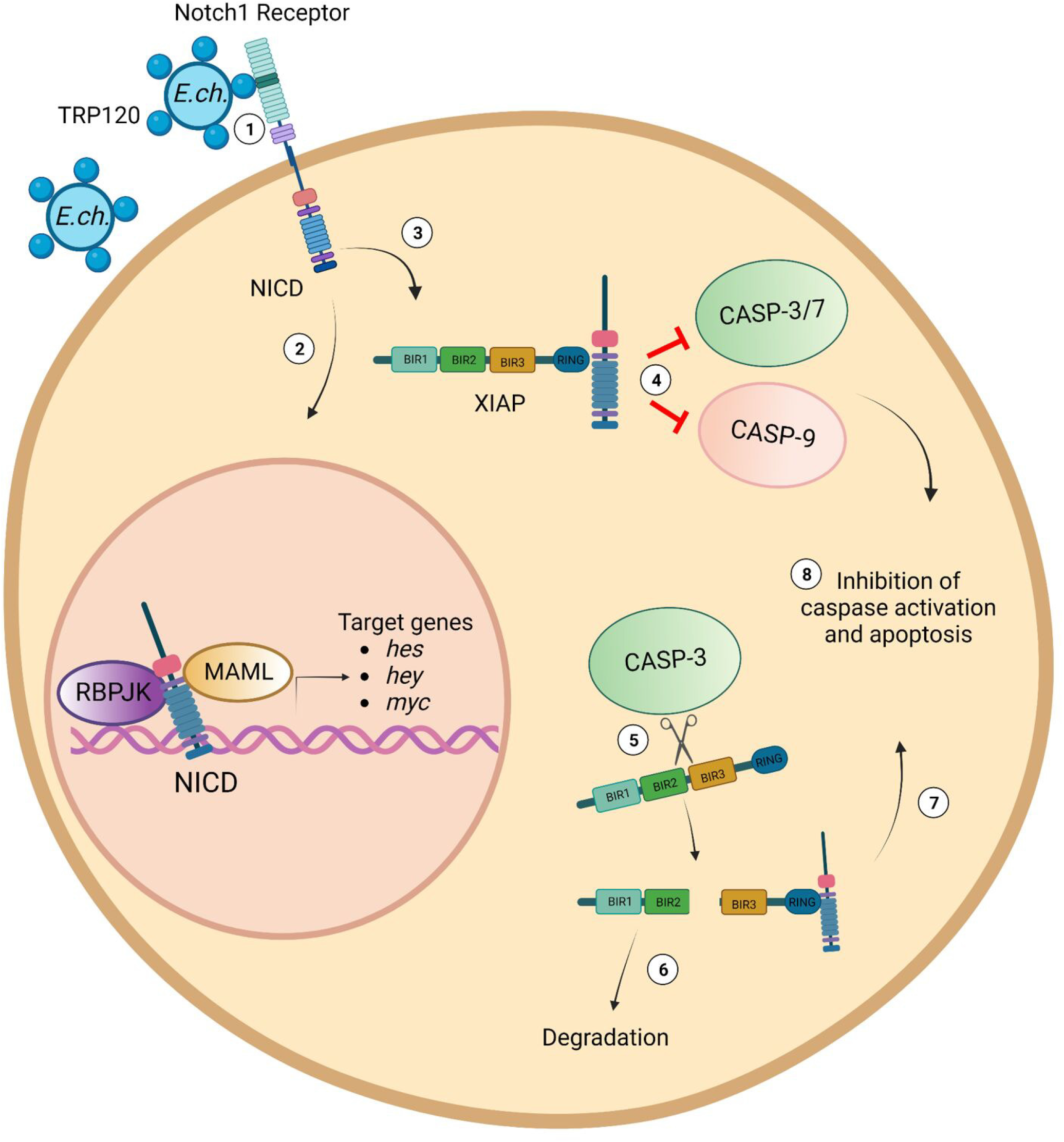
Illustration abstract of *E. chaffeensis* TRP120 Notch-mediated apoptosis inhibition. Infectious dense-cored *E. chaffeensis* with T1SS TRP120 effector on the surface binds (1) to the ligand binding region (LBR) of the Notch-1 receptor via the Notch ligand memetic motif (Patterson, Velayutham et al. 2022) and activates Notch signaling. (2) NICD translocates to the nucleus and activates Notch gene transcription and (3) binds directly to XIAP prevent XIAP degradation, (4) inhibiting Caspase-3, −7 and −9 activation. (5) Active Caspase-3 may also cleave XIAP into two fragments: BIR1 and −2 and BIR3-RING. (6) BIR1 and −2 are more rapidly degraded in comparison to BIR3-RING. (7) BIR3-RING can bind to NICD and (8) both full-length and cleaved XIAP-BIR3-RING/NICD complexes can inhibit caspase activation and apoptosis.

Pathogens have evolved various means of subverting innate immune defense of the host for survival (2, 43, 55–60). One of the most well-studied mechanisms of bacterial pathogens is targeting intracellular signal transduction cascades. Pathways such as MAPK and NF-kB are manipulated by various pathogens (39, 61–63). Manipulation of these innate immune and inflammatory pathways are regulated through bacterial effector proteins and host-pathogen interactions (3, 64). In addition, inhibition of apoptosis is well-documented as a mechanism used by bacteria and viruses to subvert innate immune defense (2, 56, 62, 65, 66). Although intracellular bacterial pathogens manipulate apoptosis by various mechanisms, exploitation of the Notch signaling pathway is a strategy that has not been reported.

The Notch signaling pathway is an evolutionary, highly conserved cell signaling pathway that is conserved in Metazoans (67, 68). The Notch pathway is known to play critical roles in cellular homeostasis, differentiation and cell proliferation (68). Recently, evidence has demonstrated Notch signaling to also play roles in innate immunity and inflammation, including regulation of Toll-like receptor expression, inflammatory cytokines, macrophage activation, MHC class II expression, B- and T-cell development, autophagy and apoptosis (37, 69–73). Interestingly, Notch signaling is activated in macrophages by LPS (74), *Mycobacterium bovis* BCG (75), and as we recently reported, *E. chaffeensis* (5). Activation of Notch by LPS and by *M. bovis* BCG was associated with the regulation of cytokine signaling through different mechanisms. The expression of canonical Notch ligand, Jagged-1, was induced by LPS in a JNK-dependent manner (74). In comparison, *M. bovis* BCG was shown to upregulate Notch-1 and activate the Notch-1 signaling pathway, leading to the expression of SOCS3, a negative regulator of cytokine signaling (75). In comparison, *E. chaffeensis* has been demonstrated to activate Notch signaling through a Notch SLiM memetic motif found within the TR domain of the TRP120 effector resulting in downregulation of TLR 2/4 (5, 13).

Interestingly, Notch has been shown to inhibit apoptosis by directly interfering with the ubiquitination of the most potent inhibitor of apoptosis, XIAP (47). NICD directly binds the BIR3-RING domain of XIAP to inhibit autoubiquitination. Inhibition of XIAP autoubiquitination results in stabilization of XIAP levels, leading to inhibition of apoptosis (47). In this study we demonstrated increases in XIAP expression over the course of *E. chaffeensis* infection. A cleavage product was observed at later time points which correlated with the XIAP BIR3-RING domain. Studies have demonstrated this domain to be a potent inhibitor of the intrinsic apoptotic pathway through direct binding to Caspase-9 (48). XIAP sequesters Caspase-9 in a monomeric state, which serves to prevent catalytic activity (76). Previous studies have demonstrated that *E. chaffeensis* inhibits apoptosis through the intrinsic apoptotic pathway by blocking the BCL-2 pathway (33). In addition, we have recently demonstrated that engagement of the BCL-2 anti-apoptotic cellular programming during *E. chaffeensis* infection is caused by activation of the Hedgehog signaling pathway (2). Induction of BCL-2 resulted in inhibition of Caspase-3 and −9, preventing activation of intrinsic apoptosis (2). Therefore, evidence supports the inhibition of intrinsic apoptosis as a survival mechanism for *E. chaffeensis.* Importantly, both rTRP120 and the TRP120 Notch SLiM ligand memetic peptide upregulated XIAP expression in a timedependent manner, with significant upregulation occurring at later time-points, as demonstrated with *E. chaffeensis* infected cells. These data support the role of the TRP120 induced Notch signaling activation leading to increased XIAP levels.

Increases in NICD and XIAP levels occurred simultaneously and temporally. We have previously determined that NICD expression increases during infection, attributed in part to *E. chaffeensis* TRP120 ubiquitinating and degrading the Notch negative regulator, FBW7 (15). Here, we demonstrate NICD to both colocalize and directly bind with XIAP at later time points of *E. chaffeensis* infection. Interestingly, colocalization of NICD and XIAP occurred in both the cytoplasm and the nucleus during *E. chaffeensis* infection. Selective localization of pro-caspases in different subcellular compartments have been previously demonstrated (77). Pro-caspase and active Caspase-3, −7 and −9 are mainly found in the cytosolic fraction; however, Caspase-3 and −9 were also found in the mitochondrial fraction, while Caspase-7 in the microsomal fraction in untreated Jurkat T lymphocytes (77). Caspase-3 was the only major caspase found in the nucleus (77). XIAP expression has been found to be mainly cytoplasmic; however, is also present in the nucleus in specific cell types. XIAP nuclear translocation have been previously associated with aberrant cell division and anchorage-independent growth (78, 79). Previous data has demonstrated increases in XIAP are not associated with stimulation of XIAP transcription by NICD. Therefore, further investigation is needed to determine the functional implications of NICD/XIAP colocalization in the nucleus during *E. chaffeensis* infection (47). Inhibition of Notch activation by DAPT, a γ-secretase inhibitor, reversed increases in XIAP levels during *E. chaffeensis* infection. DAPT inhibits Notch receptor enzymatic hydrolysis, NICD release and downstream transcriptional activation by inhibiting γ-secretase activity. Hence, inhibition of NICD is directly associated with decreases in XIAP levels.

Apoptosis is an important innate defense mechanism against microbial infection; however, various intracellular pathogens hijack apoptosis by inhibiting either extrinsic or intrinsic apoptosis through different mechanisms (2, 80–82). We demonstrated the importance of XIAP expression in inhibition of apoptosis during *E. chaffeensis* infection. siRNA knockdown of XIAP significantly reduced *E. chaffeensis* infection. This finding was associated with apoptosis, as demonstrated by Muse Count & Viability Assays and microscopy demonstrating cellular morphological hallmarks of apoptosis. siRNA treated cells showed significant cell blebbing, shrinkage of the cell and nuclear fragmentation. XIAP siRNA treated cells contained a significant reduction in morulae compared to scrambled siRNA cells. Interestingly, *A. phagocytophilum* also appears to inhibit apoptosis by preventing XIAP degradation (43). Cleaved fragments of XIAP were not detected in *A. phagocytophilum-infected* neutrophils (43), suggesting that XIAP degradation is blocked during *A. phagocytophilum* infection. In contrast, we detected an increase in XIAP cleavage product (30 kDa), which we identified as XIAP BIR3-RING. As previously stated, the XIAP BIR3-RING cleavage product has been shown to strongly inhibit intrinsic apoptosis. Differences in the presence of the cleavage product observed between *E. chaffeensis* and *A. phagocytophilum* are not well understood and need further investigation. These findings indicate that modulating IAPs to inhibit apoptosis may be a conserved mechanism utilized by various intracellular bacterial pathogens for survival.

Various studies have demonstrated XIAP as the most potent endogenous inhibitor of caspases due to weaker binding and inhibition of caspases by other IAP proteins. Interestingly, XIAP has been shown to inhibit both the executioner and intrinsic apoptotic pathways using various domains found within its structure. XIAP inhibits the executioner pathway by directly binding to Caspase-3 and −7 through the linker region between the BIR1 and BIR2 domains (51). As previously mentioned, XIAP also directly binds to Caspase-9 via the BIR3 domain (45, 46, 76). We have demonstrated levels of pro-Caspase-3, −7 and −9 temporally increase during *E. chaffeensis* infection. However, there were only minor changes in Caspase-8 levels during infection, demonstrating that mitochondrial-mediated apoptosis is the predominantly targeted for inhibition. By comparison, inhibition of Caspase-8 activation and Bid cleavage has been demonstrated in *A. phagocytophilum-infected* human neutrophils (43).

Previous data has shown that transcriptional levels of caspases do not change at earlier timepoints during *E. chaffeensis* infection (83). Our data correlates with these observations in that significant upregulation is not shown until 48 hpi. Pro-Caspase-3 protein levels increased at 24 hpi but decreased at 48 and 72 hpi. Cleavage of pro-Caspase-3 levels coincides with BIR3-RING cleavage products observed at 48 and 72 hpi. Activated Caspase-3 has previously been demonstrated to cleave XIAP (48), resulting in BIR1-2 and BIR3-RING fragments. The BIR1-2 fragment inhibits Caspase-3 and −7; however, BIR1-2 is a less potent inhibitor of apoptosis than full-length XIAP and may also be susceptible to further degradation (84). In comparison, the BIR3-RING fragment blocks activation of Caspase-9 by directly binding to and inhibiting activity (45, 46, 76). Therefore, activation of Caspase-3 may lead to XIAP BIR3-RING fragments that inhibit intrinsic apoptosis through direct Caspase-9 binding. Interestingly, similar evidence of inhibition of Caspase-3 and −9 activation during *A. phagocytophilum* infection has been reported where activation of Caspase-3 and −9 were linked to inhibition of XIAP degradation (43).

Inhibition of caspase activation by XIAP is mediated by the endogenous IAP inhibitor, SMAC/Diablo. During induction of apoptosis, SMAC/Diablo is processed and released from the mitochondria where it binds to the BIR2 and BIR3 domains of XIAP to antagonize XIAP activity (45, 50). SM-164 is a bivalent, SMAC mimetic that induces apoptosis (85). Treatment with SM-164 in the presence of TNF-α significantly reduced cell viability and ehrlichial load. An increase in caspase-positive apoptotic cells was shown with SM164/TNF-α treatment. Importantly, SM-164/TNF-α treated cells pre-treated with Caspase-9 inhibitor, Z-LEHD-FMK TFA, blocked full induction of apoptosis during *E. chaffeensis* infection. TNF-α has been demonstrated to inhibit apoptosis through both the extrinsic and intrinsic apoptotic pathways. Activation of the extrinsic pathway results in the cleavage of cytosolic BID to truncated p15 BID (tBID), which translocates to mitochondria and triggers cytochrome c release (86). Therefore, reversal of cell death by Caspase-9 inhibitor, Z-LEHD-FMK TFA, may occur through Bid activation. In addition, SM-164 treatment resulted in decreased pro-caspase and increased levels of cleaved Caspase-9 and −3 during infection. These results demonstrate direct correlation of XIAP activity and caspase inhibition during *E. chaffeensis* infection.

During *E. chaffeensis* infection, TRP120 ubiquitinates Notch negative regulator, FBW7, resulting in degradation. (15). Degradation of FBW7 is known to result in increased NICD levels (87). HeLa cells transfected with TRP120-C520S catalytic mutant displayed an increase in cleaved Caspase-3 and −9, demonstrating a direct relationship between FBW7 stabilization of NICD levels and subsequent increased XIAP expression and caspase inhibition. Collectively, this study serves to provide insight into the molecular details of how TRP120 Notch signaling leads to increased XIAP expression through direct interaction with NICD, leading to inhibition of caspase activation and apoptosis for *E. chaffeensis* survival.

There are multiple questions that remain to be answered regarding *E. chaffeensis* regulation of apoptosis. IAP proteins have been previously shown to interact with one another to form IAP-IAP complexes that inhibit apoptosis (88). Many of the IAP-IAP complexes consist of one or more of four key IAPs: c-IAP1, c-IAP2, XIAP and survivin. Whether XIAP functions in a complex with other IAPs to inhibit apoptosis during *E. chaffeensis* infection remains unknown. Evolutionarily conserved signaling pathways such as Notch and Hedgehog play key roles in regulation of apoptosis (35, 89, 90). TRP120 has been demonstrated to inhibit apoptosis by activation of Hedgehog signaling (2). Further investigation is needed to understand potential crosstalk between Notch, Hedgehog and potentially other signaling pathways that are activated during *E. chaffeensis* infection and associated with apoptosis regulation.

In conclusion, we demonstrated *E. chaffeensis* Notch activation results in an XIAP-mediated anti-apoptotic program. Our findings reveal an *E. chaffeensis* initiated, Notch signaling regulated, antiapoptotic mechanism involving inhibitor of apoptosis proteins (IAPs) that inhibits caspase activation (Fig 8). This study gives further insight into the molecular mechanisms used by obligate intracellular pathogens to exploit conserved signaling pathways to suppress innate defenses and promote infection.

## Materials and Methods

### Cell culture and *E. chaffeensis* infection

Human monocytic leukemia cells (THP-1; ATCC TIB-202) were maintained in RPMI medium (ATCC) supplemented with 2 mM l-glutamine, 10 mM HEPES, 1 mM sodium pyruvate, 4500 mg/L glucose, 1500 mg/L sodium bicarbonate, supplemented with 10% fetal bovine serum (fetal bovine serum [FBS]; Invitrogen) at 37°C in 5% CO_2_ atmosphere. *E. chaffeensis* (Arkansas strain) was propagated in THP-1 cells. Host cell-free *E. chaffeensis* was prepared as previously described (13). The purified ehrlichiae were resuspended in fresh RPMI medium and utilized as needed.

### Antibodies and reagents

Primary antibodies used in this study for immunofluorescence microscopy and immunoblot analysis include polyclonal rabbit (2042S, Cell Signaling Technology, Danvers MA) or mouse monoclonal α-XIAP (sc-55550, Santa Cruz Biotechnology, Dallas TX), rabbit monoclonal α-Caspase-3 (9662S; Cell Signaling Technology), rabbit monoclonal α-Caspase-7 (9494S; Cell Signaling Technology), rabbit monoclonal α-Caspase-8 (4790T; Cell Signaling Technology), rabbit α-Caspase-9 (9502S; Cell Signaling Technology), polyclonal rabbit α-Notch1 intracellular domain (07-1231; Millipore Sigma, Billerica, MA), rabbit α-TRP120-I1, rabbit monoclonal α-GAPDH (2118L; Cell Signaling Technology), and human monoclonal α-OMP-1 (91). Synthetic peptides used in this study were commercially generated (Genscript, Piscataway, NJ). The pharmacological inhibitors of XIAP, Notch and Caspase-9 in this study were SM-164 (56003S; Cell Signaling Technology), DAPT (GSI-IX) (S2215; Tocris Bioscience, Bristol, UK) and Z-LEHD-FMK TFA (S731303; Tocris Bioscience), respectively. Cell death inducers in this study included rhTNF-α (210-TA/CF; R&D Systems, Minneapolis MN) and Staurosporine (9953S; Cell Signaling Technology).

### Immunoblot analysis

Cells were infected or treated as indicated in text and figure legends and subsequently lysed with Triton-X 100 supplemented with protease inhibitor cocktail, Halt phosphatase and phenylmethylsulfonyl fluoride (PMSF) for 30 min, with lysing by pipetting every 10 min on ice. Lysates were cleared by centrifugation at 14,000 x *g* (4°C) for 20 min. Protein concentration of cleared lysates were determined by bicinchoninic acid assay (BCA assay). Laemelli buffer was added to lysates then boiled for 5 min at 95°C. Lysates were then subjected to SDS-PAGE and transferred to nitrocellulose membrane. Membranes were blocked using 5% nonfat milk in TBST and then exposed to α-XIAP, α-NICD, α-Casp-3, −7 or −9 or α-GAPDH antibodies overnight at 4°C. Membranes were washed thrice in Tris-buffered saline containing 1% Triton (TBST) for a total of 30 min followed by 1 h of incubation with horseradish peroxidase-conjugated anti-rabbit or anti-mouse secondary antibodies (SeraCare, Milford, MA) (diluted 1:10,000 in 1% nonfat milk in TBST). Proteins were visualized with ECL using a ChemiDoc It^2^ imager (UVP) and densitometry performed with VisionWorks software (ver. 8.1).

### Quantitative PCR

Uninfected, *E. chaffeensis*-infected or inhibitor-treated THP-1 cells were collected at 12, 24, 48 and 72 h intervals. RNeasy Mini Kit (Qiagen, Valencia, CA, USA) was used to purify RNA followed by cDNA synthesis (0.5 μg of RNA) using iScript RT kit (Bio-Rad) and qPCR was performed using the Brilliant II SYBR^®^ Green QPCR master mix (Agilent). PCR primer sequences included *XIAP* (F: 5’-GAGAAGATGACTTTTAACAGTTTGA-3’; R: 5’-TTTTTTGCTTGAAAGTAATGACTGTGT-3’), *CASP-3* (F: 5’-TGCAGCAAACCTCAGGGAAA-3’; R: 5’-AGTAACCCCTGCTTAATCGTCA-3’), *CASP-7* (F: 5’-ATTTAGGCTTGCCGAGGGAC-3’; R: 5’-ATGCTTGGCAGACAATGGAC-3’), *CASP-9* (F: 5’-GGCTGCTCCTGTTGGATGTA-3’; R: 5’-CCTTTTACCCTTGGTTTGGGC-3’), *DSB* (F: 5’-GCTGCTCCACCAATAAATGTATCCT-3’; 5’-GTTTCATTAGCCAAGAATTCCGACACT-3’) and GAPDH (F: 5’-GGAGTCCACTGGCGTCTTCAC-3’; R: 5’-GAGGCATTGCTGATGATCTTGAG-3’). Relative gene expression was calculated by determining the cycle threshold (Ct) value and normalizing to *GAPDH* as previously described (13).

### Co-immunoprecipitation

Magna ChIP™ A/G Chromatin Immunoprecipitation kit (MilliporeSigma, Burlington, MA) was used to investigate XIAP and NICD interactions during *E. chaffeensis* infection. Briefly, THP-1 cells were infected with *E. chaffeensis* (MOI 100) or left uninfected (control) for 24 h. Cells were harvested, and Co-IP was performed according to the manufacturer’s protocol. XIAP and NICD antibodies (Cell Signaling Technology) were used to determine interactions. IgG purified from normal serum was used as control antibody. Bound antigen was eluted, solubilized in 4X SDS sample loading buffer, and processed for immunoblot analysis. The membrane was probed with XIAP or NICD antibody to confirm pulldown. Co-immunoprecipitation was performed in triplicate experiments.

### Transfection

HeLa cells (1 × 10^6^) were seeded in a 60 mm culture dish 24 h prior to transfection. All proteins were expressed in a pcDNA3.1+C-6His vector. TRP120 full-length (pcDNA3.1+TRP120_FL_C-6His) and its HECT Ub ligase catalytic inactive mutant (pcDNA3.1+TRP120_C520S_C-6His) were cloned into the pcDNA3.1+C-6His vector at NheI/XbaI sites. pcDNA3.1+C-6His empty vector was used as a control. All vectors were added to Opti-MEM and Lipofectamine 3000 mixture and incubated for 20 min at 37°C. Lipofectamine/plasmid mixtures were added to HeLa cells and incubated for 4 h at 37°C. The medium was aspirated 4 h posttransfection and fresh medium was added to each plate and incubated for 24 h.

### Immunofluorescent confocal microscopy

THP-1 cells (1×10^6^) were infected with *E. chaffeensis* (MOI 100) for indicated time intervals at 37°C. Cells were collected and fixed using ice-cold 4% formaldehyde and washed with sterile 1X PBS five times for 5 min. Cell samples were permeabilized and blocked in 0.5% Triton X-100 and 2% bovine serum albumin (BSA) in PBS for 30 min. Cells were washed with sterile 1X PBS three times for 5 min and probed with XIAP, NICD or DSB antibodies for 1 h at room temperature. Cells were washed with sterile 1X PBST (0.1% Tween) three times for 5 min and probed with Alexa Fluor IgG (H+L) or Alexa Fluor IgG (H+L) for 30 min at room temperature, washed three times with sterile 1X PBST, and mounted with ProLong Gold antifade reagent with DAPI (Molecular Probes). Slides were imaged on a Zeiss LSM 880 confocal laser scanning microscopy. Mander’s correlation coefficients were generated by ImageJ software to quantify the degree of colocalization between fluorophores.

### TRP120 recombinant protein and peptide treatment

Recombinant TRP120 containing the tandem repeat (rTRP120-TR) or thioredoxin was expressed in a pBAD expression vector and purified as previously published (92–94). rTRP120-TR was dialyzed in 1X PBS and tested for bacterial endotoxins using the Limulus amebocyte lysate (LAL) test. TRP120 synthetic peptides were produced commercially (Genscript, Piscataway, NJ). THP-1 cells or primary monocytes were treated with 2 μg/ml of rTRP120-TR or TRX, or 1 μg/ml of synthetic TRP120 peptides for 0-72 h timepoints. Cells were collected post-treatment and immunoblot and qPCR analysis was performed.

### iRNA knockdown

Cells (1.0 × 10^6^) were transfected with ON-TARGETplus SMARTpool XIAP siRNA (3 μl) (Dharmacon, Lafayette, Co) using Lipofectamine 3000 (7.5 μl) (Invitrogen, Waltham, MA) according to the manufacturer’s instructions. Scrambled siRNA was utilized as a control in both uninfected and *E. chaffeensis-infected* THP-1 cells. The siRNA and Lipofectamine mixture were added to 250 μl of MEM medium (Invitrogen), incubated for 12 min at room temperature and added to cells in a 6-well plate. Knockdown was assessed by immunoblot analysis as previously described. Cells were knocked down for 24 h and infected with *E. chaffeensis* (MOI 100) for 24 h. Cells were then harvested after 24 h post-infection. Proliferative/cell death analysis was performed on all knockdown cells. Ehrlichial load was determined by qPCR of *dsb* gene as previously described (8). All siRNA knockdowns were performed with triplicate technical and biological replicates and significance was determined using a *t*-test analysis.

### Cell death analysis

THP-1 cells were untreated or treated with DMSO, SM-164 (100 nM) alone or in combination with Z-LEHD-FMK TFA (20 μM) and incubated at 37°C, 5% CO_2_ for 12 h. Z-LEHD-FMK TFA treatment was administered for 2 h prior to SM-164 treatment. Cell death was induced using TNF-α (100 ng/ml). Staurosporine (100 ng/ml) was utilized as a positive apoptosis control. Cells were infected with *E. chaffeensis* (MOI 50) following induction of cell death for 48 h. Apoptosis was analyzed utilizing various cell death assays.

1. *Trypan blue exclusion.* 20 μl of cell sample were collected and added to 20 μl of trypan blue. Samples were incubated at room temperature for 2 mins and read via the via Nexcelom Cellometer Mini (Nexcelom Bioscience LLC, Lawrence, MA, USA)
2. *Caspase-3/CPP32 Assay Kit.* Caspase-3/CPP32 Assay Kit (Colorimetric) [NBP2-54838] was utilized to assess the activity of Caspase-3 according to the manufacturers’ protocol.
3. *Hematoxylin and eosin (H&E).* H&E stain was utilized to assess cell morphological changes associated with cell death. Cells were collected and washed with 1X DPBS. Fresh RPMI mediated were added to the cell samples and fixed onto slides by cytospin (800 x *g*, 5 min). Cells were then fixed by acetone (1 min) and stained with Hematoxylin and eosin (1 min per stain). Slides were rinsed with DI water and dried prior to analysis using light microscopy. Images were taken using the Olympus cellSens software.
4. *Guava^®^ Muse^®^ Cell Analyzer.* Various Muse assays were utilized according to the manufacturers’ protocol to determine apoptosis:

a. The Muse^®^ Count & Viability Kit was used to determine cell count and viability (Part Number MCH100102) Viable cell count (cells/mL) and percentage viability of samples were determined.
b. The Muse^®^ MultiCaspase Kit (Part Number: MCH100109) was used to determine caspase activation and cellular plasma membrane permeabilization, or cell death. The percentage of live, caspase+, caspase+ and dead, total caspase+, and dead cells was determined.
c. The Muse^®^ Annexin V & Dead Cell Kit (Part Number: MCH100105) was used to determine live, early, and late apoptosis and cell death. The percentage of live, early apoptotic, late apoptotic, total apoptotic, and dead cells was determined.

### Statistical analysis

All data are represented as the means ± standard deviation (SD) of data obtained from at least three independent experiments done with triplicate biological replicates. Experiments performed with technical replicates are indicated in figure legends and the material and methods section. Analyses were performed using a two-way ANOVA or two-tailed Student’s *t*-test (GraphPad Prism 6 software, La Jolla, CA). P < 0.05 was considered statistically significant.

## Acknowledgments

We thank the University of Texas Medical Branch (UTMB) Optical Microscopy Core for assistance with confocal microscopy. This work was supported by the National Institute of Allergy and Infectious Diseases grant AI158422 to J.W.M., James W. McLaughlin Endowment and an NIH 1F31AI152424 predoctoral fellowships to L.L.P., and T32AI007526-20 Biodefense training fellowship to C.D.B.

We declare no conflict of interest.

## Figure Legends

**Figure S1: SAHM1 treatment decreases XIAP during *E. chaffeensis* Infection.** (A) Immunoblot and fold differences of XIAP expression in *E. ch*.-infected THP-1 cells with or without SAHM1 pre-treatment at 24 or 48 hpi. Bar graphs represent means ± SD. **, P < 0.01; ns = no significance. Experiments were performed in duplicates (n=2) and representative images are shown.

**Figure S2: Inhibition of XIAP enhances apoptosis in *E.ch.*-infected cells.** *E. ch.*-infected THP-1 cells (MOI 50) were pre-treated with DMSO or SM-164 alone (100nM, 12h) or SM164 (100nM, 12h) in combination with caspase-9 inhibitor, Z-LEHD-FMK TFA (20μM, 2h). Cell death was stimulated with TNF-α (100 ng/ml) or staurosporine (100 ng/ml, positive apoptosis control) in the indicated samples. (A) Cell viability was determined by trypan blue exclusion on the indicated treated cells. Quantification of live vs. dead cells were analyzed by Brightfield cell counting using the automated Cellometer Mini. Brightfield images were taken from the counts. Live cells are shown in green and dead cells are shown in red. (B) Percentages of live, early apoptotic, late apoptotic, total apoptotic, and dead cells were determined by the Muse^®^ Annexin V & Dead Cell Kit. Bar graphs represent means ± SD. ****, P < 0.0001. Experiments were performed in triplicate (n=3) and representative images are shown.

**Fig. S3: Pro-Caspase transcript levels are increased during *E. chaffeensis* infection**. Change in pro-Caspase-3, −7 and −9 transcription in *E.ch.* infected and uninfected THP-1 cells (0 hpi), as measured by RT–qPCR analysis. Bar graphs represent means ± SD. ****, P < 0.0001. Experiments were performed in triplicate (n=3) and representative images are shown.

